# Quantitative differences in anaerobic growth and metabolism of three different *E. coli* strains using ^13^C-Metabolic Flux Analysis

**DOI:** 10.1101/2024.03.27.586947

**Authors:** Piyush Pachauri, Trunil Desai, Ahmad Ahmad, Shireesh Srivastava

## Abstract

*Escherichia coli* are model organisms for biotechnological research owing to their abilities to grow on defined media and consume a variety of carbon sources, as well as a vast knowledge base and tools available for their manipulation. Several *E. coli* strains and their derivatives are available. However, the quantitative metabolic differences between the strains could be better characterized. In this study, we investigated the differences in the intracellular flux distribution between three commonly used strains of *E. coli* (K-12 MG1655, C, and W) using ^13^C-Metabolic flux analysis (^13^C-MFA), the gold standard in fluxomics. The labeling data was analyzed using FluxPyt, our previously developed Python-based tool. Threefold differences in the growth rates of these strains were observed when growing anaerobically on glucose in M9 medium, with *E. coli* W being the fastest (μ = 0.62 ± 0.01 h^-1^) and *E. coli* C the slowest (μ = 0.22 ± 0.01 h^-1^). The strain’s growth rates correlate with glucose uptake and flux through the oxidative pentose phosphate pathway. Interestingly, succinate was a major product in *E. coli* C only. ^13^C-MFA revealed that under these conditions, Entner–Doudoroff (ED) pathway was active in *E. coli* C and *E. coli* W but not in *E. coli* K-12 MG1655. In *E. coli* K-12 MG1655 and *E. coli* C, the Tricarboxylic Acid (TCA) cycle ran incompletely. Also, the glyoxylate shunt was inactive in all three strains. Growth, xylose intake rates, and product profiles on xylose mirrored those of glucose. Thus, this study has highlighted significant differences in the growth and metabolism of the *E. coli* strains on glucose and xylose. This will form a basis for identifying the best strain for a particular application.

## Introduction

In the realm of microbiology, *Escherichia coli (E. coli)* stands as a quintessential model organism, a microscopic workhorse that transcends its humble origins in the gut microbiota to become a linchpin in both research and industrial applications (Carreón-Rodríguez et al., 2023; Geurtsen et al., 2022; Hiller & Metallo, 2013; Khankal et al., 2008). The versatility of *E. coli* strains emerges as a critical asset for various industrial sectors. Different strains of *E. coli* have become invaluable tools in biotech endeavors, each harboring unique genetic characteristics that can be harnessed for specific purposes (Shiloach et al., 2010; Studier et al., 2009). From protein production to the synthesis of biofuels and pharmaceuticals, these strains serve as bioengineered workhorses, efficiently carrying out intricate metabolic processes (Atsumi & Liao, 2008; Bokinsky et al., 2011; Marisch et al., 2013). A wide range of E. coli strains (engineered and wild-type) available could be used as hosts to produce a commercial compound (Shiloach et al., 2010). Therefore, selecting the appropriate host strain is a critical step in the design of a bioprocess or metabolic engineering (Rumbold et al., 2009).

Understanding integrated metabolic pathways, pathway flux, and flux control lies at the core of metabolic engineering (Stephanopoulos, 1999). ^13^C-Metabolic flux analysis (^13^C-MFA) has become a gold standard in flux analysis because it enables the precise quantification of metabolic fluxes in the central metabolic pathways of an organism (Antoniewicz, 2015; Crown & Antoniewicz, 2013; Marx et al., 1996). Indeed, ^13^C-MFA is an invaluable method for the metabolic characterization of different strains or physiological states of an organism (Wiechert et al., 2001). In stationary ^13^C-MFA, ^13^C-labeled substrates are fed to the cells grown in controlled conditions. After the cells have reached an isotopic and metabolic steady state, culture is harvested, and ^13^C-labeling of proteinogenic amino acids is measured by mass spectrometry or nuclear magnetic resonance (Antoniewicz et al., 2007; Sauer et al., 1999; Zamboni et al., 2009). The intracellular fluxes are estimated by minimizing the difference between labeling data and the labeling patterns simulated by the metabolic network model based on the elementary metabolite unit (EMU) framework (Antoniewicz, 2018). ^13^C-MFA of variants of a single *E. coli* strain under different growth conditions or gene KOs have been performed previously (Arifin et al., 2014; Chen et al., 2011; Gonzalez et al., 2017a; Long & Antoniewicz, 2019b). However, to our knowledge, no prior studies have compared internal flux distributions of different wild-type *E. coli* strains in standard, identical growth conditions.

In the present study, we compare the growth and metabolic fluxes of three wild-type *E. coli* strains, viz., *E. coli* K-12 MG1655 (hereafter, *E. coli* MG1655), *E. coli* C, and *E. coli* W, in anaerobic growth in a minimal medium using ^13^C-MFA to gain a comprehensive understanding of their metabolic capabilities and differences. We have also compared these strains’ growth and fermentation profiles in M9 minimal medium with xylose as a carbon source. *E. coli* MG1655 is a commonly used wild-type strain for studying biochemical genetics (Blattner et al., 1997). *E. coli* W is a fast-growing strain that can use sucrose as a carbon source (Archer et al., 2011). *E. coli* W has been used as a host strain for producing ethanol and penicillin G acyclase (PGA) (Ohta et al., 1991; Sobotková et al., 1996). *E. coli* C is a relatively less explored strain in metabolic engineering studies, but it has been considered a relatively good host for bacteriophages (Monk et al., 2016). All these strains belong to Biosafety level 1, making them suitable for industrial applications.

Our study has identified significant differences in these strains’ growth rates, product profiles, and intracellular fluxes. These findings could be helpful when selecting an *E. coli* strain for specific applications, such as producing a target compound from lignocellulose biomass. The ^13^C-MFA data of these strains can also be used as a foundation for designing metabolic engineering strategies.

## Material and methods

### Materials

Media and chemicals were purchased from Sigma-Aldrich Chemicals Private Limited (Bangalore). Isotopic-labeled glucose was purchased from Omicron Biochemicals, Inc. (IN, USA): [1-^13^C] glucose (98–99%) and [U-^13^C] glucose (99%). The isotopic purity of the labeled glucose was validated by gas chromatography-mass spectrometry (GC-MS) (Long et al., 2016) M9 minimal medium was used for all experiments.

### Strains and culture conditions

*E. coli* MG1655 was obtained from Dr. Yazdani, Microbial Engineering Group, ICGEB, New Delhi, while *E. coli* W (DSMZ 1116) and *E. coli* C (DSMZ 4860) were purchased from Leibniz Institute DSMZ (Germany). M9 medium with glucose (3 g/L) or xylose (10 g/L) as the sole carbon source was used for culturing. Glucose and xylose cultures were incubated at 37 °C and 150 rpm.

### Culture with ^13^C -labeled glucose for ^13^C-MFA

A single colony of each strain was first inoculated into 50 mL M9 medium with 3g/L glucose taken in 100 mL serum bottles and grown overnight at 37 °C and 150 rpm. This primary culture was inoculated into 50 mL fresh M9 medium in 100 mL serum bottles containing a mixture of ^13^C -labeled glucose (80% mol/mol [1-^13^C] glucose and 20% mol/mol [U-^13^C] glucose, 3.0 g/L in total) (Chen et al., 2011; Hua et al., 2006). The bottles were sealed with rubber stoppers to prepare anaerobic cultures, and grade-I nitrogen gas was flushed into the headspace of the bottles through a sterile needle for 30 minutes. The nitrogen gas was sterilized before entering the bottle by passing it through an air filter (MDI, 0.22 μm). Another needle was inserted into the rubber stopper to allow venting. The second needle was removed before the first needle to prevent any contamination from the air. (Monk et al., 2016). The primary culture was inoculated into the above anaerobic culture medium with an optical density at 600 nm (OD_600)_ of 0.04 using a sterile syringe and needle. Cultures were maintained at neutral pH (6.9 to 7.1) by intermittently adding sterile 1 M KOH using a syringe and a needle (the total volume of 1 M KOH used was less than 5 mL). Cell growth was monitored by measuring OD_600_ using a spectrophotometer (Ultrospec 3100 pro, Amersham Biosciences). Samples for OD_600_ and extracellular metabolite measurements were also drawn using a syringe and a needle. Cell pellets were collected at an OD_600_ of 0.4 ± 0.02 for isotopic labeling analysis of amino acids using GC-MS. Additional samples were taken after ∼30 minutes to confirm the isotopic steady state. Labeling patterns in these additional samples were compared with that of experimental samples. This labeling pattern was within the standard deviations of the experimental samples.

### Culture with xylose

A single colony of each strain was first inoculated into 50 mL M9 medium with 10 g/L xylose taken in 100 mL serum bottles and grown overnight at 37 °C and 150 rpm. This primary culture was inoculated into 50 mL fresh M9 medium in 100 mL serum bottles with 10 g/L xylose with an initial OD_600_ ≈ 0.04. Bottles were sealed with rubber stoppers, but nitrogen was not flushed into the headspace. So, the oxygen in the headspace was available for the cells (semi-anaerobic growth). The cultures were grown at 37 °C and 150 rpm, and pH (6.9-7.1) was maintained by adding sterile 1 M KOH.

### Measurement of extracellular metabolites

Extracellular metabolites (glucose, xylose, ethanol, acetate, succinate, and lactate) were measured using High Performance Liquid Chromatography (HPLC). 1 mL culture sample was drawn at regular intervals and centrifuged at 10000 rpm for 3 minutes at 4 °C. The supernatants were stored at -20 °C until further analysis using HPLC. 10 μL of supernatant was injected via an autosampler in an HPLC machine (Agilent) with an Aminex HPX 87H (300*×*7:8 mm) column. The column temperature was maintained at 40 °C. The mobile phase (4 mM H_2_SO_4_) was passed through the column at a 0.3 ml/min flow rate.

### Calculation of extracellular fluxes (substrate uptake rate/ product secretion rates)

Extracellular fluxes were calculated from the regression analysis of extracellular metabolite measurements at different time points. First, the amount of substrate consumed/product formed per biomass generated was determined by plotting the substrate/product concentration (mmol/L) versus biomass concentration (g-DW/L) and quantifying the slope (Y) from regression analysis. Then, extracellular fluxes are calculated using the following equation (Long & Antoniewicz, 2019a)

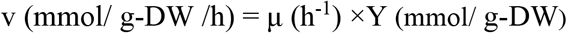

where v is the specific substrate uptake or product secretion rate (flux) and μ is the specific growth rate of the culture. OD_600_-to-biomass concentration conversion factor was determined to convert OD_600_ values to grams of dry weight per liter (gDW/L) biomass concentrations, and this factor was determined to be 0.37 ± 0.03 g-DW/L OD_600_.

### GCMS method for amino acid detection

#### Hydrolysis of cells

Cells (1.5 mL of OD_600_ = 0.4 ± 0.02) were centrifuged (10000 rpm, 4 °C, 3 minutes). The supernatant was discarded, and the pellet was resuspended in 6 M HCl. The suspension was baked at 105 °C for 24 hours in a dry bath to hydrolyze the cells. The hydrolysate was dried at 95 °C in a dry bath. The dried hydrolysate was stored at -20 °C.

### Derivatization of amino acids

The hydrolysates were dried again at 105 °C for 15 minutes before derivatization. They were then resuspended in dimethyl formamide (DMF, 20 μl) and transferred to a fresh tube. 20 μl N-tertbutyldimethylsilyl-N-methyltrifluoroacetamide with 1% (wt/wt) tertbutyldimethyl-chlorosilane (TBDMSTFA)was added, and the solutions were incubated at 85 °C for 1 hour. The derivatized amino acid solutions were centrifuged (10000 rpm, 3 min, 4 °C) to separate the debris. The supernatants were transferred into vials for GC-MS analysis (Zamboni et al., 2009).

The derivatized amino acid samples were analyzed using GC-MS with an Agilent 7890A GC system equipped with an HP-5column (30 m *×* 0.25 mm ID, 0.25 μm film thickness, Varian) coupled with Agilent 7000 QQQ MS. 1 μl of the sample was injected in the split mode 10:1. The signals were recorded in full scan mode (m/z 50 to 650, 250 scan/ms). An electron ionization system with an ionization energy of 70 eV was used, and 99.99% pure Helium gas was used as the carrier gas at a constant flow rate of 1.1 mL/min. The temperatures of the mass transfer line and the injector were set at 220 °C and 250 °C, respectively. The oven temperature was programmed as follows: the initial temperature was set to be 100 °C for 2 min, then increased by 5 °C/min up to 180 °C and again increased by 10 °C/min up to 300 °C and was held for 0.5 min. The amino acid fragments were identified, and their retention times were recorded by comparing the fragmentation patterns with the NIST mass spectral library using AMDIS and Mass Hunter software.

### Calculation of mass isotopomer distribution (MID)

The extracted ion chromatogram of each mass fragment was exported to a CSV file. Using this data and the retention times of the amino acids, the chromatograms were integrated using a bespoke Python code. The measurement standard deviations for all MIDs were 0.02 (Zamboni et al., 2009). Although the actual standard deviations in the measurements were lower than 0.02, the deviation value considers the inherent errors in the GC-MS machine.

### Calculation of flux distribution

The metabolic network of *E. coli* was constructed based on a previously published metabolic network (Gonzalez et al., 2017b). The coefficients of amino acids in the biomass reactions were taken from a genome-scale metabolic model of *E. coli* (Orth & Palsson, 2012). Flux estimation and confidence interval calculations were done using the FluxPyt package (Desai & Srivastava, 2018) in Python (version 3.6.) FluxPyt package is freely available at Sourceforge (https://sourceforge.net/projects/fluxpyt/). The measured flux values constrained the metabolic model (Table 1). This narrows down the feasible solution space and allows a more accurate estimation of flux distribution. The metabolic network had 28 free flux parameters, of which one reaction, i.e., the glucose uptake rate, was assigned a fixed value.

**Table 1:**
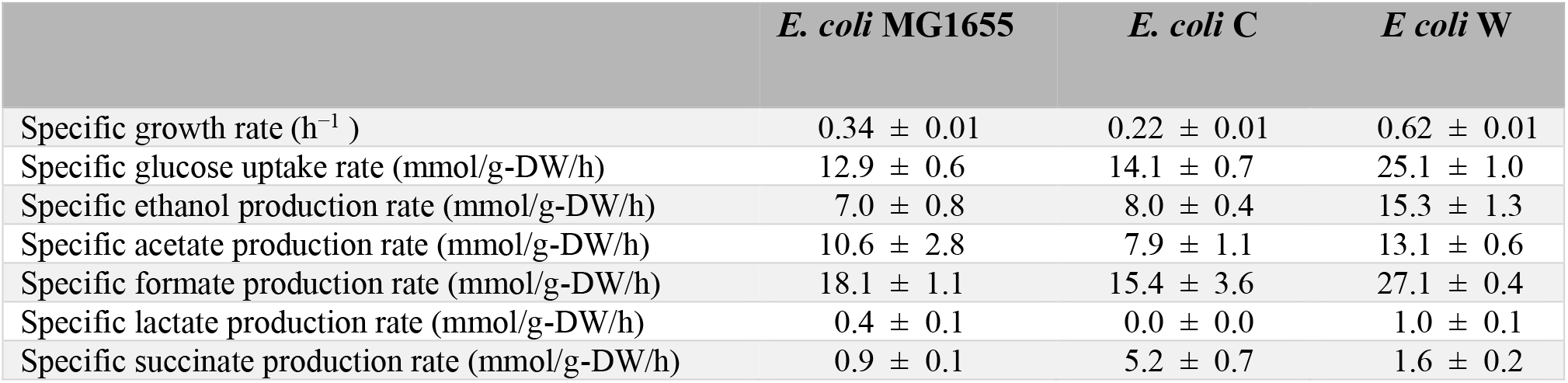
Specific growth rates and extracellular fluxes (specific rates) under anaerobic growth on glucose. All data represent mean ± standard deviation obtained from three independent cultures.

For MFA models, 76 MID data points were fitted to 28 free flux parameters. Hence, the degree of freedom of the system was 48. 10 iterations of non-linear least squares fitting were performed on the models to estimate the fluxes using random initial values of free flux parameters. The following formula calculated the sum of squared residues (SSR):

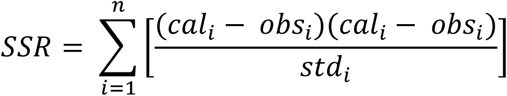

Where *n* is the number of MID measurements. *cal*_*i*_, *obs*_*i*_, and *std*_*i*_ are the *i*^*th*^ calculated MID, observed MID, and standard deviation, respectively.

### Goodness of fit analysis

The ^13^C-MFA fitting results were subjected to the χ^2^ - test. Assuming the model is correct, and data are without gross measurement errors, the minimized SSR is a stochastic variable with a χ2-distribution. The number of degrees of freedom equals the number of fitted measurements *n* minus the number of estimated parameters *p*. The acceptable range of the SSR is between χ^2^_α/2_(n-p) to χ^2^_1-α/2_(n-p), where α is the certain chosen threshold value, for example, α = 0.05 for 95% confidence interval (Antoniewicz et al., 2006).

## Results

### Growth parameters and uptake/ secretion rates with glucose as carbon source

Table 1 shows the measured specific growth rates, specific glucose uptake rates, and specific production rates (extracellular fluxes). The growth rates of the *E. coli* strains varied widely. *E. coli* W was the fastest-growing strain, with a growth rate of 0.62 ± 0.01 h^−1^, whereas *E. coli* C was the slowest-growing strain (growth rate of 0.22 ± 0.01 h^−1^). The growth rate of *E. coli* MG1655 (0.34 ± 0.01 h^−1^) and extracellular fluxes were consistent with the previous reports (Chen et al., 2011; Fischer et al., 2004; Hua et al., 2006). Figure 1(A-C) shows the growth and fermentation profiles of the strains.

**Figure. 1.**
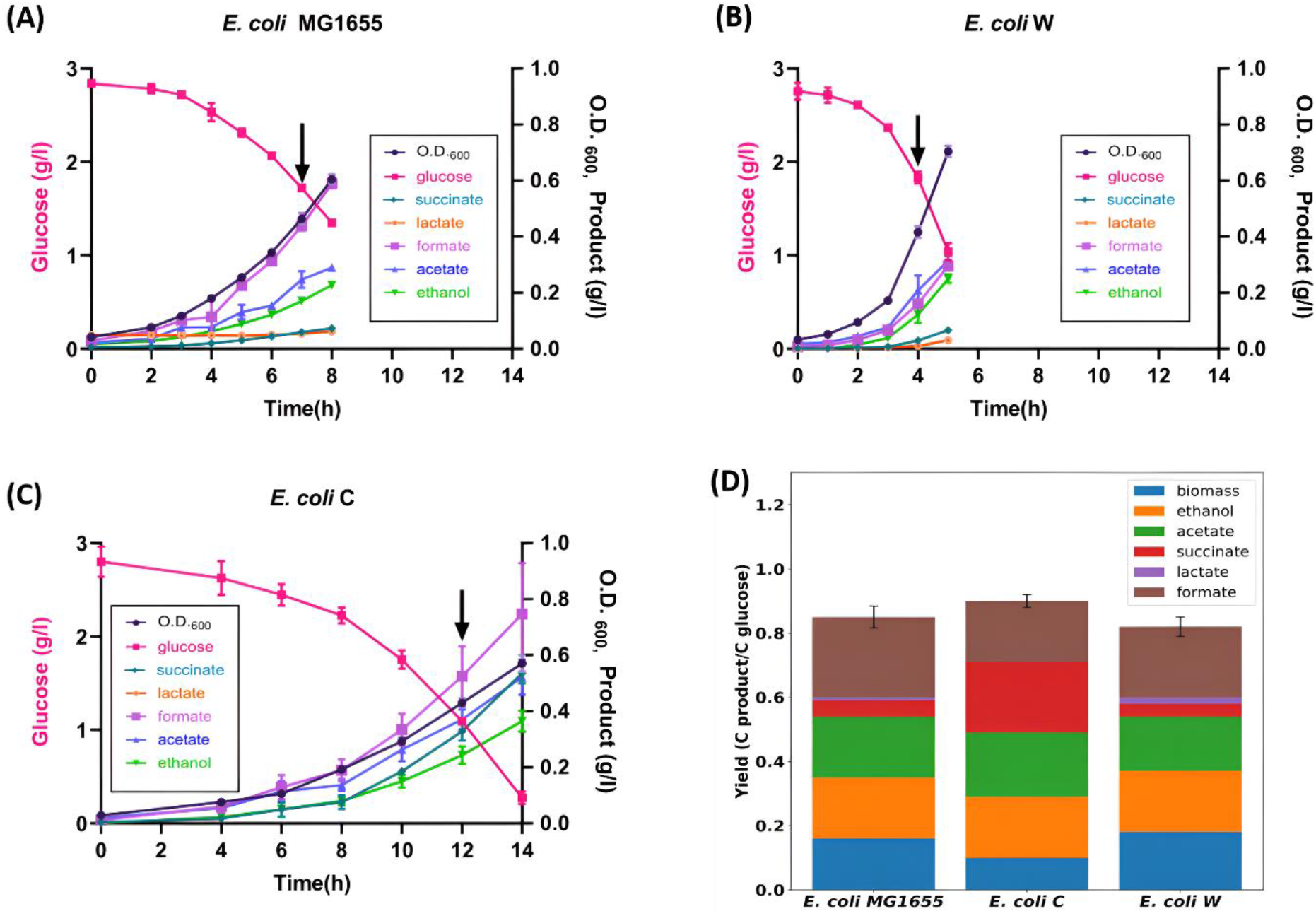
(A), (B) and (C) represent the fermentation and growth profiles of *E. coli* MG1655, *E. coli* W, and *E. coli* C, respectively, with glucose as a carbon source. (D) Comparison of different strain’s carbon yield. Yields are calculated in terms of glucose uptake rate. All data represent mean ± standard deviation obtained from three independent cultures. Black arrows indicate the sampling time points for the measurements of ^13^C-labeling of proteinogenic amino acids.

The specific glucose uptake rate was measured to be highest in *E. coli* W (25.1 ± 1.0 mmol/g-DW/h), whereas in *E. coli* C and *E. coli* MG1655, the measured specific glucose uptake rates were similar (14.1 ± 0.7 mmol/g-DW/h, 12.9 ± 0.6 mmol/g-DW/h, respectively).

The fermentation product profiles of the strains show some unique differences and similarities among strains. The major fermentation products were formate, acetate, and ethanol in all three strains, while lactate and succinate were also detected in small amounts.

*E. coli* W also had the best product formation rates, producing ethanol with the highest specific production rate (15.3 ± 1.3 mmol/g-DW/h). This rate was around double the measured rates in *E. coli* C and *E. coli* MG1655 (8.0 ± 0.4 mmol/g-DW/h and 7.0 ± 0.8 mmol/g-DW/h, respectively). Acetate and formate production rates were also found to be highest in *E. coli* W. Interestingly, despite having the slowest growth, *E. coli* C had the highest succinate production rate of 5.2 ± 0.7 mmol/g-DW/h.

Figure 1(D) compares the carbon yields of different strains. Yields are calculated in terms of glucose uptake. Carbon dioxide was not measured. Overall, the by-product profiles differed across the strains. In the case of *E. coli* C, succinate yield was around 4 times higher than the other two strains, and its biomass yield was lowest. Lactate was detected in the case of *E. coli* MG1655 and *E. coli* W. We captured about 80-90% of the carbon yields in the by-products measured. This is similar to the values reported by other researchers (Gonzalez et al., 2017a; Monk et al., 2016)).

### ^13^C -Metabolic flux analysis

^13^C-MFA was carried out to investigate the similarities and differences in central carbon metabolism of the *E. coli* strains. The best-fit solutions for all three strains had SSR values within the acceptable χ^2^ range (30.75 to 69.02). The SSR for *E. coli* W, *E. coli* C, and *E. coli* MG1655 were 56.82, 66.61, and 32.62, respectively. Figure 2 shows the estimated flux maps for the three strains. Figure 3 shows the estimated flux values together with 68% and 95% confidence intervals for six representative metabolic fluxes in the network model: glycolysis (Embden–Meyerhof pathway, EMP, R03), oxidative pentose phosphate pathway (oxPPP, R10), phosphoenolpyruvate carboxylase (PPC, R28), pyruvate formate lyase (PFL, R20), citrate synthase (CS, R23), and succinate dehydrogenase (SDH, R26). The complete flux results are provided in the supplementary table. The estimated flux distributions of the three strains showed differences. The central carbon metabolic pathways, including glycolysis, oxPPP, and TCA cycle, were active in all strains. Most of the glucose was metabolized via glycolysis and oxidative PPP in all three strains. In the case of *E. coli* MG1655, around 70% of the glucose was metabolized via glycolysis, while the remaining 30% was processed in oxPPP. The Entner-Doudoroff (ED) pathway, glyoxylate shunt, and malic enzyme were inactive in *E. coli* MG1655, as reported by previous studies (Chen et al., 2011; Crown et al., 2015). The TCA cycle was found to be running incompletely as no significant flux was observed through α-ketoglutarate (AKG) to succinyl-CoA.

**Figure 2.**
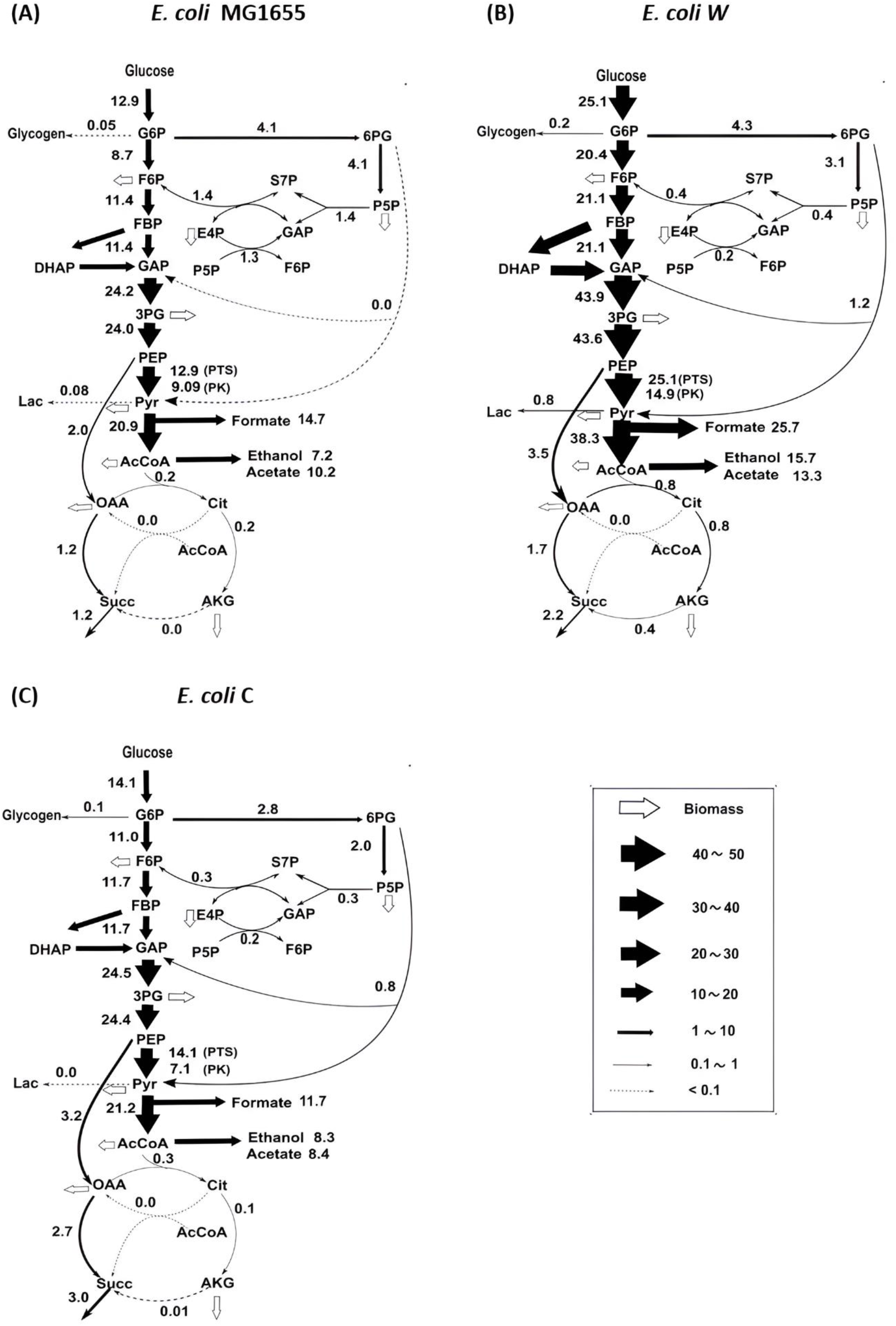
Metabolic flux maps of three different *E. coli* strains grown in batch culture in anaerobic conditions with glucose as carbon source. Fluxes were determined using ^13^C -MFA. Flux values (in mmol/gDW/h) represent the optimal solution. The line width reflects the flux rate. Complete flux results and confidence intervals are provided in Supplemental materials.

**Figure. 3.**
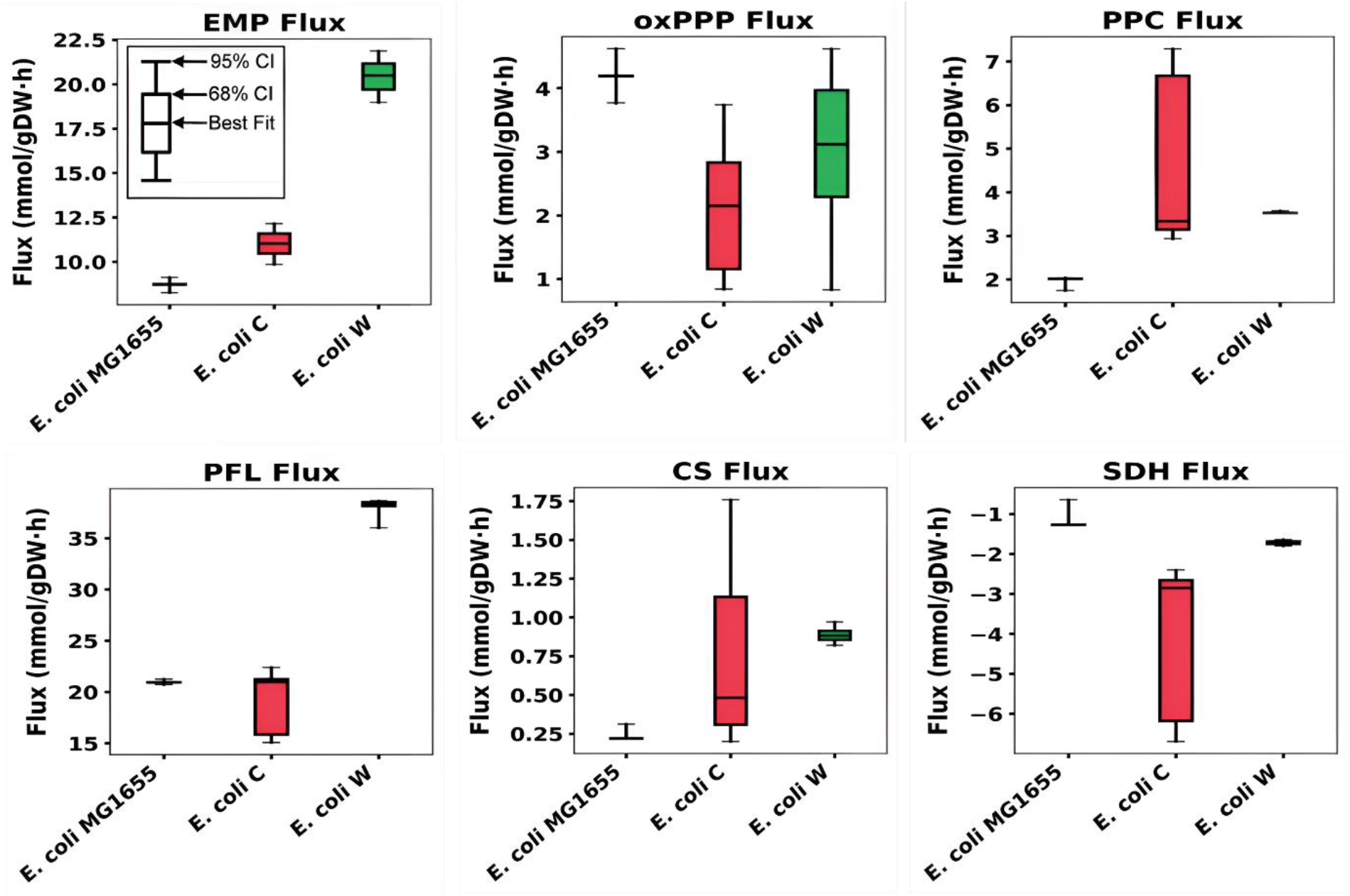
Comparison of estimated metabolic fluxes of *E. coli* K-12 MG1655, *E. coli* C and *E. coli* W. The 68% and 95% confidence intervals are shown for six representative metabolic fluxes in the model: glycolysis (Embden–Meyerhof pathway, EMP, R03), oxidative pentose phosphate pathway (oxPPP, R10), phosphoenolpyruvate carboxylase (PPC, R28), pyruvate formate lyase (PFL, R20), citrate synthase (CS, R23), and succinate dehydrogenase (SDH, R26).

In *E. coli* W, due to the high glucose uptake rate, higher absolute values of fluxes were found in the glycolytic pathway with 83% flux via glycolysis, while only about 17% of the glucose uptake flux was metabolized through oxPPP. The glyoxylate shunt and malic enzyme were inactive. Interestingly, a significant flux (about 5%) was observed in the ED pathway, driving carbon flux from oxPPP to the glycolytic pathway. Unlike *E. coli* MG1655, the TCA cycle was running completely here.

*E. coli* C was metabolizing 80% of the glucose flux through the glycolytic route, and the rest, 20%, was going into oxPPP. The glyoxylate shunt pathway and malic enzyme were inactive, similar to the observation in the other two strains. However, the ED pathway was active. The TCA cycle was found to be bifurcated because of insignificant flux from AKG to succinyl-CoA. Interestingly, the highest flux of succinate was observed in *E. coli* C, which was produced through the reductive branch of the TCA cycle.

### Xylose fermentation under microaerobic conditions

Our initial experiments showed that *E. coli* C could not grow on xylose in the minimal medium under anaerobic conditions (data not shown). Therefore, the growth on xylose was compared under microaerobic conditions, i.e., the cultures were not bubbled with N_2_ gas as done for the glucose cultivation. Table 2 shows the measured specific growth rates, specific xylose uptake rates, and specific production rates (extracellular fluxes).

**Table 2:** Specific growth rates and extracellular fluxes. All data represent mean ± standard deviation obtained from three independent cultures.

|  | *E. coli* MG1655 | *E. coli* C | *E. coli* W |
| --- | --- | --- | --- |
| Specific growth rate (h<sup>-1</sup>) | 0.38 ± 0.01 | 0.27 ± 0.02 | 0.62 ± 0.01 |
| Specific xylose uptake rate (mmol/g-DW/h) | 11.8 ± 2.3 | 9.5 ± 0.07 | 15.8 ± 0.9 |
| Specific ethanol production rate (mmol/g-DW/h) | 5.8 ± 1.1 | 4.6 ± 0.6 | 5.0 ± 0.3 |
| Specific acetate production rate (mmol/g-DW/h) | 7.1 ± 0.6 | 7.7 ± 1.4 | 7.7 ± 0.9 |
| Specific formate production rate (mmol/g-DW/h) | 11.5 ± 0.9 | 9.7 ± 1.8 | 13.1 ± 1.5 |
| Specific lactate production rate (mmol/g-DW/h) | 0.2 ± 0.1 | 0.7 ± 0.2 | 0.5 ± 0.1 |
| Specific succinate production rate (mmol/g-DW/h) | 0.8 ± 0.1 | 1.7 ± 0.4 | 0.8 ± 0.2 |

As observed in the case of glucose, with xylose also, *E. coli* W outperformed the other two strains in terms of growth rate (0.62 ± 0.01 h^−1^) and specific xylose uptake rate (15.8 ± 0.9 mmol/g-DW/h) while *E. coli* C was the slowest, with a 0.27 ± 0.02 h^−1^ growth rate and the lowest specific xylose uptake rate (9.5 ± 0.07 mmol/g-DW/h). Figure 4 (A-C) shows growth and fermentation profiles of these strains with xylose in M9 medium.

**Figure 4.**
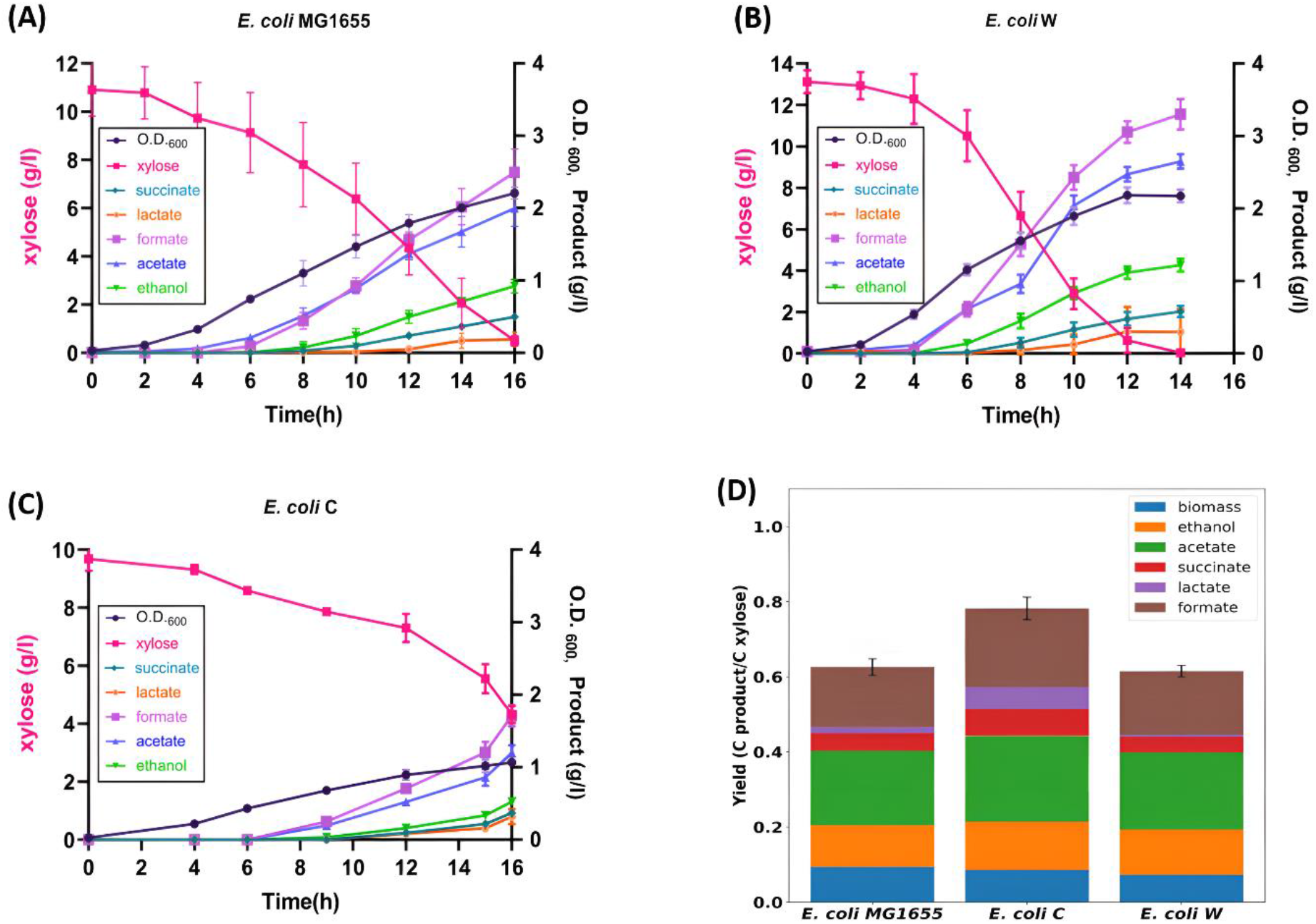
(A), (B) and (C) represent the fermentation and growth profiles of *E. coli* MG1655, *E. coli* W, and *E. coli* C, respectively, with xylose as a carbon source. (D) Comparison of different strain’s carbon yield. Yields are calculated in terms of xylose uptake rate. All data represent mean ± standard deviation obtained from four independent cultures.

In terms of fermentation products, all three *E. coli* strains produced ethanol, acetate, and formate as major products, while lactate and succinate were also produced, although in small amounts. This observation was in agreement with the previously done studies (Gonzalez et al., 2017a).

*E. coli* W was the best producer of formate, with a specific production rate of 13.1 ± 1.5 mmol/g-DW/h. However, the specific production rates for acetate and ethanol did not show much variation among the strains. *E. coli* C outperformed the other two strains in succinate production with about two times higher specific production rate than the other two strains (1.7 ± 0.4 mmol/g-DW/h).

The fermentation products and biomass could capture approximately 60% of the carbon in the case of *E. coli* W and *E. coli* MG1655 as shown in Figure 4(D). However, in the case of *E. coli* C, 78% of total carbon was recovered in terms of by-products and biomass. These results suggest that a significant amount of carbon from xylose was released as CO_2_, as observed earlier by other groups (Gonzalez et al., 2017a). The C-yields of formate, acetate, and ethanol did not show much variation among the strains. All three strains also produced Succinate and lactate, although in small amounts. *E. coli* C performed better among all three strains regarding succinate and lactate yields.

## Discussion

This study identifies significant differences in the central carbon metabolism of three different strains of *E. coli*, which is an industrially important microbial species. These strains have been reported to have comparatively similar genomes and metabolic reaction networks (Baumler et al., 2011; Monk et al., 2013; Vieira et al., 2011). Our findings demonstrate that these *E. coli* strains differ significantly in terms of growth and fermentation profiles, even though their genomic structures are comparable.

The growth profiles of the strains on glucose (Figure 1A-C) show significant variation, with about a 3-times difference between the maximum and minimum growth rates. Some studies have reported higher growth rates for these strains with glucose under anaerobic conditions; however, they have used some additional medium components like Wolf’s vitamin solution other than just M9 minimal medium components (Monk et al., 2016).

The product profiles suggest that *E. coli* W should be the strain of choice for ethanol production from glucose and xylose as it has the highest uptake rates for these sugars compared to the other two strains tested. *E. coli* C outperformed the other two strains in terms of succinate production despite having a relatively low glucose uptake rate. Succinic acid has been identified by the US Department of Energy as one of the top 12 biomass-derived building block chemicals (Liu et al., 2017).

^13^C-MFA proved effective in elucidating the intracellular flux distribution of the strains. The choice of labeled glucose used can impact the values and the precision of the fluxes calculated. In this study, we have used the isotopic tracer mixture (4:1) of [1-^13^C] glucose and [U-^13^C] glucose. Crown et al. 2015, conducted 14 parallel labeling experiments and found that the isotopic tracer mixture (4:1) of [1-^13^C] glucose and [U-^13^C] glucose performed well for glycolysis, the oxidative pentose phosphate pathway, and the ED pathway. However, its performance was poor for the TCA cycle and anaplerotic fluxes. Under anaerobic conditions, previous studies suggest minimal flux through the TCA cycle. Therefore, we used this specific tracer mixture to evaluate these strains’ central carbon metabolism variations accurately. ^13^C-MFA results show that most of the carbon is metabolized by the glycolytic pathway, oxPPP, and fermentation reactions, forming ethanol, acetate, and formate. The TCA cycle reactions were found to be operating at low fluxes, mainly to produce biomass precursors. In the case of *E. coli* MG1655 and *E. coli* C, the TCA cycle was bifurcated, as observed by previous reports (Chen et al., 2011). A consistent finding of ^13^C-MFA was that the reducing branch of the TCA cycle was carrying more flux due to the net production of oxaloacetic acid (OAA) from phosphoenolpyruvate (PEP), and the glyoxylate shunt was inactive. The ED pathway was active in *E. coli* W and *E. coli* C, another significant observation. The ED pathway has been reported to have lower costs for protein and ATP yields than the glycolytic route for carbon metabolism (Chen et al., 2016). *E. coli* has been engineered to overexpress the ED pathway to produce commercially valuable compounds like isopropyl alcohol (Okahashi et al., 2017).

The growth rate of *E. coli* W is roughly twice that of *E. coli* MG1655, and it can be noted that the glucose uptake rate of *E. coli* W is about twice that of *E. coli* MG1655. However, ^13^C-MFA data shows that the relative flux in oxPPP is higher in *E. coli* MG1655 than in *E. coli* W. These observations suggest that the growth rate of *E. coli* MG1655 is limited by the amount of flux in the glycolytic pathway rather than in oxPPP.

Although the glucose uptake rates of *E. coli* MG1655 and *E. coli* C are similar, *E. coli* C grows approximately 35% slower than *E. coli* MG1655. The ^13^C-MFA data shows that the flux in oxPPP in *E. coli* C is approximately 30% lower than in *E. coli* MG1655. Thus, we hypothesize that the low flux in oxPPP could limit its growth rate.

Finally, the xylose fermentation study shows that these strains can utilize xylose as a carbon source and produce compounds of industrial importance. ^13^C-MFA of the strains with xylose was not performed because of the high cost of labeled xylose. The observed growth rates indicate that strains that grew faster on glucose also performed better on xylose. Also, the fermentation product profiles on xylose follow a similar pattern to those observed with glucose.

## Acknowledgement

Author contributions: PP, TD and AA designed and conducted the experiments, analyzed the data and wrote the manuscript. SS designed and supervised the research as well as reviewed the manuscript.

PP acknowledges research fellowship from Department of Biotechnology (DBT), Ministry of Science and Technology, India. TD acknowledges research fellowship from Council for Scientific and Industrial Research (CSIR), India.

